# Ready-to-eat foods: A potential vehicle for spread of coagulase-positive staphylococci and antibiotic-resistant *Staphylococcus aureus* in Buea municipality, South West Cameroon

**DOI:** 10.1101/2021.09.30.462650

**Authors:** Seraphine Nkie Esemu, Sally Tabe Njoh, Lucy M. Ndip, Nene Kaah Keneh, Jerome Achah Kfusi, Achiangia Patrick Njukeng

## Abstract

**Background:** The consumption of ready-to-eat (RTE) foods contaminated with coagulase-positive staphylococci (CoPS) and especially *Staphylococcus aureus* puts consumers at potential risk of foodborne disease or colonization and subsequent infection. This cross-sectional study determined the levels of CoPS and the presence of *S. aureus* in RTE foods sold in Buea municipality.

**Method:** A total of 420 RTE food samples comprising 70 each of cake, bread, fruit salad, meat-hot-pot, suya and boiled rice were randomly purchased from February to August 2020. The CoPS counts were determined by culturing on Baird-Parker agar and *S. aureus* identified by amplification of the *nuc* gene using polymerase chain reaction. All *S. aureus* isolates were screened for the presence of classical staphylococcal enterotoxin genes and each isolate challenged with 11 antibiotics to determine their antibiotic susceptibility profiles. Oxacillin-resistant *S. aureus* were analyzed for the presence of *mec*A gene.

**Result:** Overall, 161 (38.3%) samples had detectable levels of coagulase-positive staphylococci ranging from 2.0-5.81 log_10_CFU/g. Based on CoPS levels, 37 (8.81%) of the 420 RTE food samples, only fruit salad and meat-hot-pot, had unsatisfactory microbiological quality. A total of 72 *S. aureus* isolates, comprising 52.78% from fruit salad, 16.67% from meat-hot-pot, 12.5% from boiled rice, 9.72% from suya, 5.56% from bread and 4.17% from cake were recovered. None of the *S. aureus* isolates possessed any of the classical enterotoxin genes. All the isolates were sensitive to vancomycin and ofloxacin while 68 (94.44%) and 66 (91.67%) were sensitive to oxacillin and ciprofloxacin, respectively. Resistance to penicillin (93.06%) was highest followed by amoxicillin (91.67%) and erythromycin (79.17%). Four isolates were identified as methicillin-resistant *S. aureus* all of which carried the *mec*A gene. A total of 24 antibiotypes were identified.

**Conclusion:** Our findings showed that RTE foods sold in the Buea municipality are likely vehicles for transmission of CoPS and antibiotic-resistant *S. aureus*.

## 1. Introduction

Ready-to-eat (RTE) foods are perishable food types that are ready for consumption, without any further heat treatment or washing, most often at the point of sale [1]. Convenience, ease of production, affordability, palatability, availability and unique flavours are some of the appealing factors that make RTE foods very popular and cherished by people of all age groups and particularly workers in urban areas, students and road side dwellers [2-4]. The popularity of RTE foods varies from country to country and depends on the food type and the staple diets of the population [5]. In low- and middle-income countries like Cameroon, RTE vendors have been associated with low literacy level leading to lack of knowledge about good hygiene and food handling practices [3, 6]. Furthermore, some RTE foods are prepared, stored and served under unhygienic conditions and vending is often done in outdoor environments, thereby exposing the foods to aerosols, insects, and rodents, which may serve as sources of food contaminants [7, 8].

Food safety in the community is a major focus area in public health (Gizaw, 2019). Three methods, considered gold standards, have been recommended for the determination of the microbiological quality and safety of RTE foods. These methods are aerobic colony count, identification of hygiene indicator organisms and identification of specific food borne pathogens (Table 1).

**Table 1:**
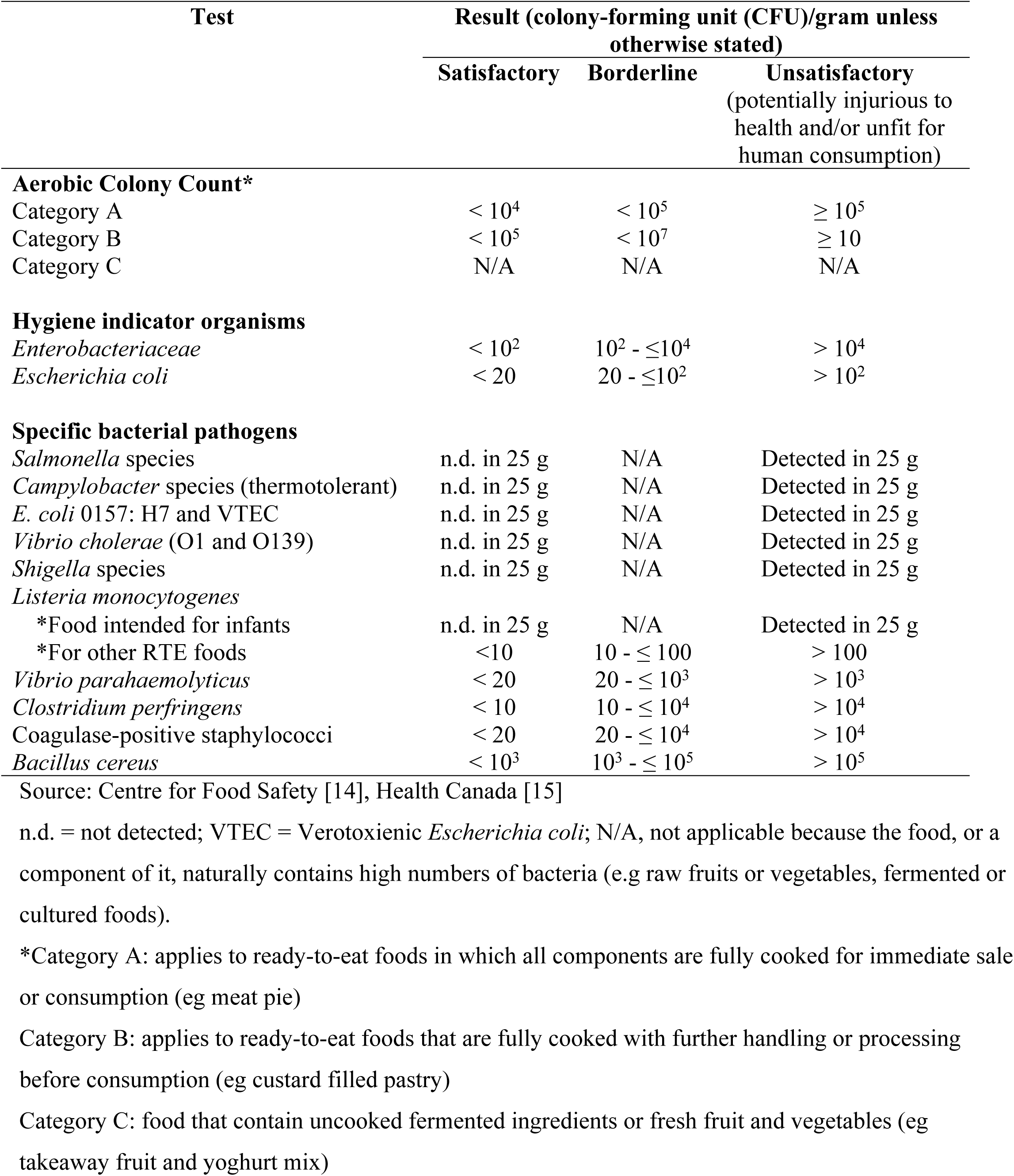
Microbiological guidelines on the interpretation of results for ready-to-eat foods

Although these methods can be used in combination, each of them is sufficient to provide evidence for the microbiological quality and safety of RTE foods [10]. Despite having many advantages, RTE foods continue to be associated with a growing number of foodborne illnesses and outbreaks mainly due to bacterial contamination [11, 12]. So far, coagulase-positive staphylococci are considered the main bacterial etiology of food poisoning [13].

Coagulase-positive staphylococci (CoPS) are opportunistic pathogens that can exist as commensals in humans, companion and food-producing animals but can cause severe or even lethal diseases [16]. They are facultative anaerobic Gram-positive, non-spore forming spherical-shaped bacteria. At least nine species of CoPS have been identified and they include *Staphylococcus aureus, S. hyicus, S. intermedius, S. pseudintermiedius, S. lutrae, S. schleiferi* subsp. *coagulans, S. delphini, S. argenteus* and *S. schweitzeri* [16, 17). Although several of these *Staphylococcus* species can produce staphylococcal enterotoxins (SEs), the majority of staphylococcal food poisoning is attributed to SE produced by *S. aureus* [18, 19]. The number of coagulase-positive staphylococci per gram or per milliliter of RTE food sample can be used to determine the microbiological quality and safety of RTE foods [20].

*Staphylococcus aureus* is one of the major bacterial agents causing foodborne disease in humans worldwide [21]. Some pathogenic strains of this bacterium elaborate heat-stable SEs and so far, 23 SEs have been identified comprising five major classical types (SEs: SEA to SEE) and the non-classical SE-like toxins (SEl: SEG to SEU) [13, 22]. Approximately 95% of food poisoning outbreaks are caused by the classical enterotoxins [23]. Ingestion of SEs containing food can induce severe symptoms, including vomiting and high fever with or without nausea and diarrhea, with rapid onset in typically less than 8 h (usually between 3 and 4 h) [24]. Consumption of RTE food contaminated with *S. aureus* has been reported in several developed countries including U.S.A., Hong Kong, Germany, Japan and Italy [1]. In China, several RTE food products have been associated with the introduction of microbiological hazards, including *Listeria monocytogenes, Cronobacter, Salmonella* and *S. aureus* [25]. Contamination by toxigenic *S. aureus* in RTE food is a major public health challenge in both developing and developed countries and the prevalence of this bacterium in RTE foods needs to be monitored and controlled [12, 26].

Over the years, several pathogenic strains of *S. aureus* isolated from RTE food have developed resistance to one or more commonly used antibiotics used in treatment of infection. Antibiotic-resistant bacteria (ARB) are a major threat to global public health. In order to effectively counter the threat of infections with antibiotic-resistant bacteria, it is critical to identify potential ways that humans can be exposed to these bacteria [27]. Although foods have been reported as vehicles for the transmission of antibiotic-resistant bacteria, the role of RTE foods in the spread of these pathogens is scantily documented [25]. Some RTE foods have been reported to carry antibiotic-resistant enterococci [5], antibiotic-resistant *Staphylococcus* species [28] and other bacterial species [29, 30]. Unfortunately, the presence of antibiotic-resistant bacteria in RTE food is not routinely investigated in low- and middle-income countries and data are only available from a small number of studies [23]. Hence, this study was carried out to investigate the levels of CoPS, determine the prevalence and characteristics of *S. aureus* isolated from commonly consumed RTE food samples collected in Buea, South West Cameroon.

## 2. Materials and methods

### 2.1 Study area

This study was carried out on RTE foods from vendors in Buea municipality; the administrative headquarter of the south west region of Cameroon. Buea is located on the slope of mount Cameroon between latitude 4°14″ north of the equator and longitude 9°20″ east of the Greenwich Meridian with an altitude of about 1000m above sea level [31]. Buea is a cosmopolitan and multicultural town and had an estimated population of 300,000 inhabitants in 2013 [32]. This town is the seat of many educational institutions at the primary, secondary and tertiary levels and is experiencing a very high rate of population growth and urbanization. At least 7000 people relocate to Buea each year according to the municipal council statistics [33].

### 2.2 Study design and RTE food types

This was a cross-sectional study carried out from February to August 2020. Buea was purposifuly selected because of its large concentration of vendors with intensive food vending activities throughout the week, from Monday through Sunday and from early hours of the day to late evening. These vendors are patronized by people-from-all-walks-of-life particularly shoppers, workers, passers-by and students.

Six types of RTE foods (fruit salad, suya (roasted beef), cake, boiled rice, bread and meat-hot-pot (cooked meat in tomato sauce) were included in this study. These RTE food types were chosen based on their popularity and round-the-clock availability. They were purchased randomly from street food vendors and only one sample was collected per vendor.

### 2.3 Collection of food samples

A total of 420 RTE food samples, comprising 70 of each food type, were purchased randomly and periodically from street vendors. Upon purchase, each sample was aseptically put in a sterile zip lock bag, assigned a unique code, and placed in a cool box containing ice packs with temperature maintained at +2 to +6°C. Samples were transported to the Laboratory for Emerging Infectious Diseases, University of Buea within an hour of collection for processing.

### 2.4 Enumeration of coagulase-positive staphylococci

The surface colony count technique was used for the enumeration of coagulase-positive staphylococci in each sample as described previously [20]. Briefly, a 500 µL inoculum from a 10-1 dilution (in Peptone saline diluent) of each RTE food sample homogenate was spread on the surface of Baird-Parker agar medium and plate allowed to stand for 15 min to allow the inoculum to be absorbed into the surface of the agar. Each sample was inoculated in duplicate and the inoculation was done within 45 min of preparation of the sample homogenate. The culture plate was incubated at 37 °C for 24-48 h. At the end of incubation, plates were examined for typical colonies of coagulase-positive staphylococci (grey-black, shiny and convex with a diameter of 1-2.5 mm surrounded by a dull halo). Only plates with colony counts of between 10 and 300 colonies per plate were considered for enumeration.

#### 2.4.1 Confirmation of coagulase-positive staphylococci

Five colonies of each type (or all colonies if less than five) were subcultured for confirmatory testing for CoPS based on DNase production and tube coagulase positivity. Briefly, a DNase agar plate was divided into five segments and each segment was spot inoculated with a different colony. Plates were incubated at 37 °C for 24 h. DNase production was evident by a defined zone of clearing surrounding the inoculated spot after flooding the DNase plate with 1N HCl and decanting excess HCl after about 30 sec. Results were read within 5 min following the application of HCl. Tube coagulase test was performed by aseptically transferring 500 µL of human plasma to a sterile round bottom tube and inoculating the plasma with an equal volume of the suspension of the colony to be tested. Incubation was done at 37 °C and the tubes examined for clotting every 30 min for 4 h by carefully tilting the tubes. If negative after 4 h, the tube was re-examined at 24 h.

The number of coagulase positive staphylococci per g of sample was calculated as follows:

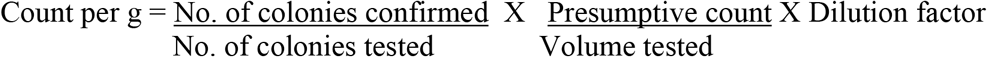

### 2.5 Molecular identification of *Staphylococcus aureus*

All colonies that were used for the confirmation of CoPS were further investigated for the identification of *S. aureus* by PCR targeting the species-specific thermonuclease (*nuc*) gene of *S. aureus*. DNA was extracted from each colony using the simple boiling method. Briefly, 150µL of phosphate buffered saline (PBS) were pipetted into a sterile 1.5 mL Eppendorf tube and a loop ful of bacterial colony suspended in the PBS. The tubes were vortexed at 14000 rpm for 10 sec. The bacterial solution was heated in a water bath at 100°C for 15min after which it was removed and immediately chilled in ice for another 15min. The bacterial solution was allowed to thaw at 37 °C in the water bath and again chilled in ice for another 15min. The tubes were centrifuged at full speed (14000rpm) for 5 min and the supernatant was pipetted into another labeled tube and used as template DNA in PCR assay.

Presumptive *S. aureus* isolates were confirmed by amplification of a 280 bp fragment of the *nuc* gene of *S. aureus* using primer sequences previously described (Table 2).

**Table 2:**
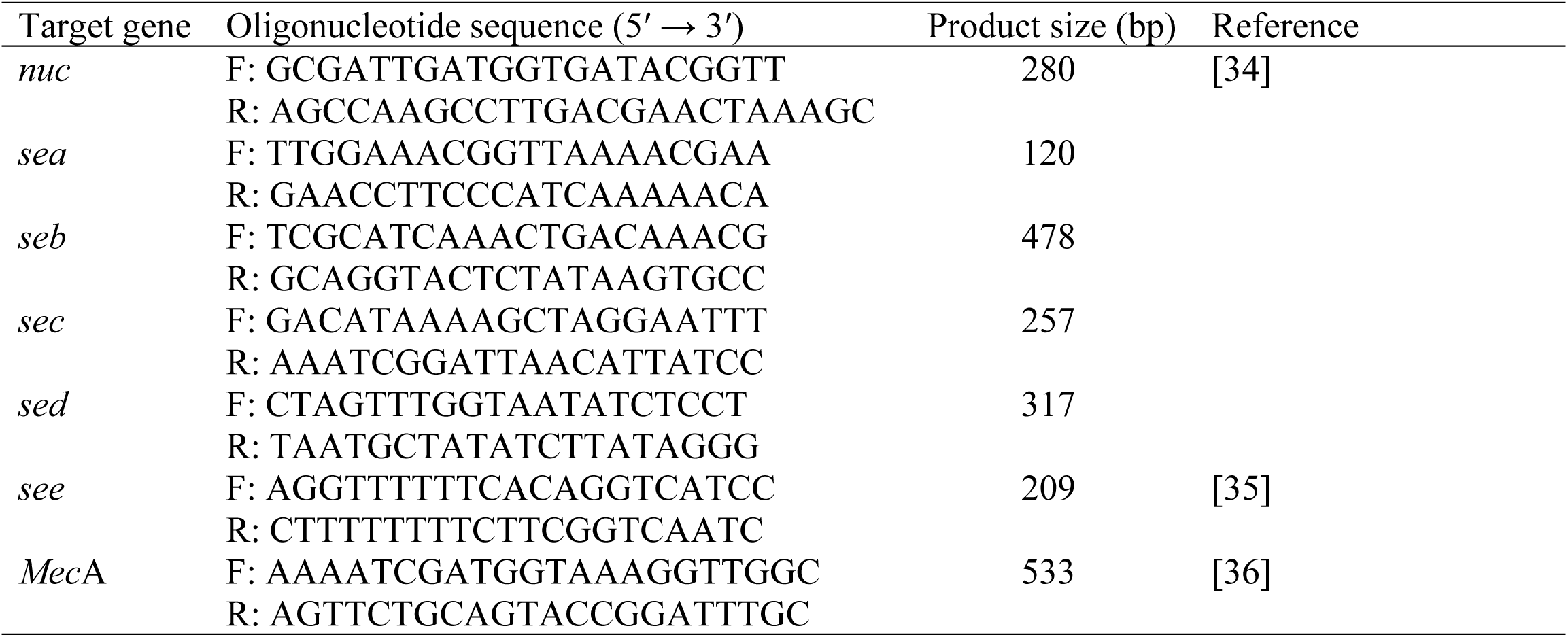
Primer sequences of *S. aureus* used for PCR assay in this study

Each PCR reaction in this study comprised 2X BioMix Red master-mix (10μL), 1μL of each primer from a 10μM working stock (final concentration, 0.5μM), 1μL DNA template and nuclease free water to make a final volume of 20µL. Each PCR run had a negative control which was nuclease-free water and a positive control which was a previously characterized *S. aureus* isolate stored in the laboratory.

The PCR cycling conditions were optimized with initial denaturation at 95°C for 5min followed by 35 cycles of denaturation at 94°C / 60sec, annealing at 54°C / 40sec and extension at 72°C / 60sec. Final extension was at 72°C for 7min and the reaction was stopped by holding tubes at 4°C until removed from the thermal cycler (MyCycler™ Thermal Cycler BIORAD, USA). The PCR products were electrophoresed in a 1.5% agarose gel stained with SYBR Safe DNA Gel stain (Invitrogen) for 1 hour at 90V in 1X Tris borate EDTA, visualized and photographed with a Molecular Imager Gel Doc XR system (BIO-RAD, Hercules, CA, USA).

### 2.6 Detection of classical enterotoxin genes in *S. aureus*

All confirmed coagulase-positive *S. aureus* isolates were screened for the presence of **S**. *aureus* enterotoxins types A, B, C and D using multiplex PCR [35] with primers previously described (Table 1). The multiplex PCR run was optimized at initial denaturation for 5 min at 94 °C followed by 30 cycles of denaturation at 94°C for 60sec, annealing at 50°C for 60sec and extension at 72°C for 60sec. A final extension at 72°C for 5min was done at the end of the cycles. A singleplex PCR run was performed for detection of enterotoxin E using the same cycling conditions except for the annealing temperature that was set at 48°C. All PCR products were electrophoretically separated in 1.5% agarose gel as described above.

### 2.7 Detection of *mec*A gene in methicillin-resistant *S. aureus* isolates

*Staphylococcus aureus* isolates that were phenotypically confirmed as methicillin-resistant were further screened by PCR for the presence of the *mec*A gene. This gene encodes a modified penicillin-binding protein designated as PBP2a with reduced affinity for β-lactams. The amplification of the 533bp fragment of the *mec*A gene was optimized under the following conditions: initial denaturation at 95°C for 5min, 35 cycles of denaturation at 94°C for 1min, annealing at 50°C for 1min, extension at 72°C for 1min with a final extension at 72°C for 5min, and cooling to 4°C. Similarly, PCR products were separated electrophoretically as described above.

### 2.8 Antibiotic susceptibility testing of *S. aureus* isolates

Antibiotic susceptibility testing (AST) was performed using the Kirby-Bauer disk diffusion method on Mueller–Hinton agar (Oxoid, UK) according to the guidelines of the Clinical Laboratory Standards Institute [37]. Each *S. aureus* isolate was challenged with a panel of 11 antibiotics viz: vancomycin (VA-30µg), chloramphenicol (C-30µg), gentamicin (CN-30µg), erythromycin (E-15µg), clindamycin (DA-2µg), ciprofloxacin (CIP-5µg), penicillin (P-10 I.U), ofloxacin (OFX-5µg), amoxicillin (AML-30µg), azithromycin (AZM-30µg) and oxacillin (OX-1µg). Antibiotics were selected to represent different antibiotic classes and also comprised commonly used and most available antibiotics for the treatment of staphylococcal-related infections in human and veterinary medicine.

Pure colonies (two to five) of each *S. aureus* isolate from nutrient agar plate were used to prepare the inoculum for AST. The turbidity of the inoculum was adjusted to that of 0.5 MacFarland standard and used to inoculate the Mueller-Hinton agar using the spreading method. Antibiotic disks were applied firmly on the agar surface and plates incubated at 35°C for 24h. The diameter of the zone of inhibition was measured and interpreted as resistant, susceptible and intermediate according to CLSI [37] guidelines. *Staphylococcus aureus* isolates that showed resistance to antibiotics in three or more antibiotic classes were considered multidrug-resistant.

### 2.9 Data quality assurance, management and statistical analysis

The handling and analysis of samples and bacterial isolates were done following standard operating procedures to ensure the quality and reliability of study findings. The sterility of culture media was maintained and confirmed by the incubation of uninoculated culture media as negative control. Data were entered into Microsoft Excel 2010, exported into SPSS V27.0.1.0 and analyzed using descriptive statistics such as prevalence, percentages, mean and standard error. Coagulase-positive staphylococci counts were transformed to log_10_CFU/mL prior to statistical analysis. Data on the prevalence of *S. aureus* and multidrug resistant isolates were analysed using a chi-squared test to determine whether there were significant differences in their prevalence between sample types. The confidence level was held at 95% and *p*-value at less than 5% for all analysis.

## 3. Results

### 3.1 Coagulase-positive staphylococci counts

The CoPS counts in the RTE foods ranged from 2.0-5.81 log_10_CFU/g. Overall, 161 (38.3%) of the RTE food samples had detectable levels of CoPS contamination. The fruit salad samples were the most contaminated (51, 72.9%) while suya samples were the least contaminated (12, 17.1%) (Table 3). A total of 652 bacterial isolates were screened in the confirmatory testing for CoPS.

**Table 3:** Coagulase-positive staphylococci counts obtained from the ready-to-eat foods analyzed

| RTE food type | Number with CoPS (%) | CoPS counts (log<sub>10</sub>CFU/g) |  | Mean±SE |
| --- | --- | --- | --- | --- |
|  |  | Minimum | Maximum |  |
| Suya (n=70) | 12 (17.1) | 2.0 | 2.91 | 2.60±0.07 |
| Fruit salad (n=70) | 51 (72.9) | 2.38 | 5.81 | 4.35±0.10 |
| Boiled rice (n=70) | 17 (24.3) | 2.3 | 3.91 | 3.40±0.13 |
| Bread (n=70) | 27 (38.6) | 2.0 | 3.95 | 2.88±0.11 |
| Meat-hot-pot (n=70) | 33 (47.1) | 2.48 | 4.94 | 3.77±0.11 |
| Cake (n=70) | 21 (30.0) | 2.23 | 3.92 | 2.83±0.09 |
| **Total (n=420)** | **161 (38.3)** |  |  |  |

RTE, ready-to-eat; CoPS, coagulase-positive staphylococci; CFU, colony forming unit; SE, standard error.

Based on CoPS counts, only 37 (8.81 %) of the 420 RTE food samples investigated and comprising only of fruit salad and meat-hot-pot, had unsatisfactory or unacceptable quality. More than half (259, 61.67%) of the RTE food samples had satisfactory quality (Table 4).

**Table 4:** Microbiological quality of the RTE food types based on coagulase-positive staphylococci counts

| RTE food type | Microbiological quality based on CoPS count |  |  |
| --- | --- | --- | --- |
| | Satisfactory [$<1.3$<br>$\log_{10}\text{CFU/g}$] (%) | Borderline [$1.3 \leq 4$<br>$\log_{10}\text{CFU/g}$] (%) | Unsatisfactory [$>4$<br>$\log_{10}\text{CFU/g}$] (%) |
| Suya (n=70) | 58 (82.86) | 12 (17.14) | 0 (0.0) |
| Fruit salad (n=70) | 19 (27.14) | 23 (32.86) | 28 (40.0) |
| Boiled rice (n=70) | 53 (75.71) | 17 (24.29) | 0 (0.0) |
| Bread (n=70) | 43 (61.43) | 27 (38.57) | 0 (0.0) |
| Meat-hot-pot (n=70) | 37 (52.86) | 24 (34.29) | 9 (12.86) |
| Cake (n=70) | 49 (70.0) | 21 (30.0) | 0 (0.0) |
| **Total (n=420)** | **259 (61.67)** | **124 (29.52)** | **37 (8.81)** |

RTE, ready-to-eat; CoPS, coagulase-positive staphylococci; CFU, colony forming unit

### 3.2 Prevalence of *Staphylococcus aureus* in food samples

Molecular confirmation of *S. aureus* relied on the amplification of the *nuc* gene by singleplex PCR. The amplified PCR products showed the desired band of required size (280 bp) when separated on a 1.5 % (W/V) agarose gel (Fig1).

**Fig 1:**
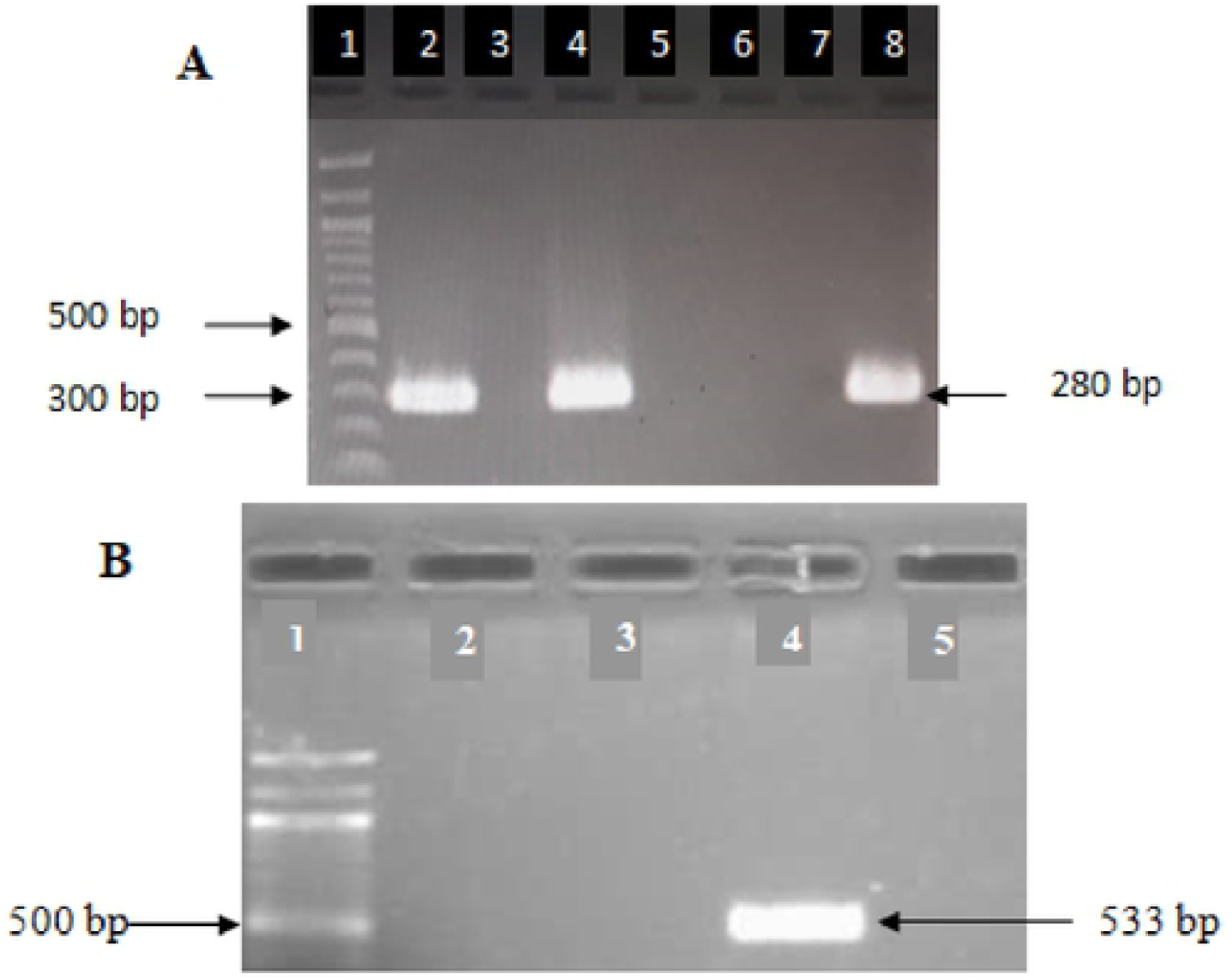
Electrophoretic separation of PCR products on 1.5% agarose gel for detection of *S. aureus* genes: (A) Amplification of *nuc* gene. Lane 1, 100 bp molecular weight marker. Lane 2, positive control. Lanes 3, 5, 6 and 7 are negative samples. Lanes 4 and 8 are positive samples. (B) Amplification of *mec*A gene. Lane 1, 100bp molecular weight marker. Lanes 2, 3 and 5 are negative samples. Lane 4 shows positive sample.

Out of the 652 CoPS isolates screened, 72 (17.14 %) were confirmed to possess the *nuc* gene and confirmed to be *S. aureus*. The 72 *S. aureus* isolates comprised 52.78% from fruit salad, 16.67% from meat-hot-pot, 12.5% from boiled rice, 9.72% from suya, 5.56% from bread and 4.17% from cake were recovered from 46 RTE food samples. Overall, *S. aureus* was detected in 10.95 % of all RTE food samples examined with some samples having more than one *S. aureus* isolates. Similarly, fruit salad was the most contaminated sample with *S. aureus* followed by boiled rice and meat-hot-pot (Table 5).

**Table 5:** Prevalence of *Staphylococcus aureus* among the ready-to-eat food samples

| RTE food type | Number of RTE food with |  |  |
| --- | --- | --- | --- |
|  | *S. aureus* (%) | Methicillin-resistant *S. aureus* (%)* | Enterotoxigenic *S. aureus* (%) |
| Suya (n=70) | 5 (7.14) | 0 (0.0) | 0 (0.0) |
| Fruit salad (n=70) | 21 (30.0) | 1 (3.33) | 0 (0.0) |
| Boiled rice (n=70) | 8 (11.43) | 0 (0.0) | 0 (0.0) |
| Bread (n=70) | 3 (4.29) | 1 (33.33) | 0 (0.0) |
| Meat-hot-pot (n=70) | 7 (10.0) | 2 (28.57) | 0 (0.0) |
| Cake (n=70) | 2 (2.86) | 0 (0.0) | 0 (0.0) |
| **Total (n=420)** | **46 (10.95)** | **4 (8.69)** | **0 (0.0)** |

\*Percentage calculated on the number of RTE food with *S. aureus*

The *mec*A gene (Fig 1) was detected in 5.56 % (4/72) of the *S. aureus* isolates from three RTE food types while none of the *S. aureus* isolates possessed any of the classical enterotoxin genes (Table 5).

### 3.3 Antibiotic susceptibility of *Staphylococcus aureus* isolates

All the *S. aureus* isolates were sensitive to vancomycin (100 %) and ofloxacin (100 %) while 68 (94.44 %) and 66 (91.67 %) of the bacterial isolates were sensitive to oxacillin and ciprofloxacin, respectively. On the other hand, the isolates showed high resistance to penicillin (93.06 %) followed by amoxixillin (91.67 %) and erythromycin (79.17 %) (Table 6). Four isolates were identified as methicillin-resistant *S. aureus* (MRSA) based on their resistance to oxacillin. Each of the MRSA isolates was resistant to at least five antibiotics. Of the 72 *S. aureus* isolates, 45 (62.5 %) were multidrug resistant. In this study, multidrug resistance was defined as resistance to one antibiotic in three or more classes of antibiotics.

**Table 6:**
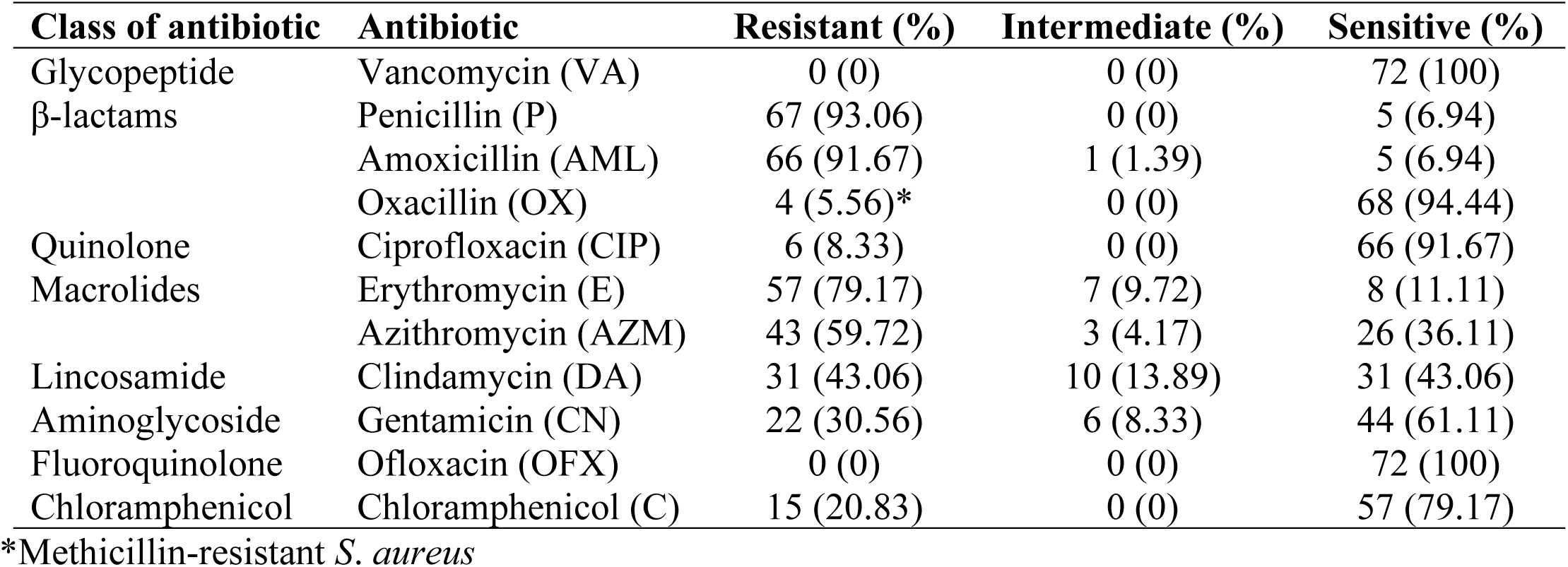
Susceptibility of *S. aureus* isolates to the different antibiotics tested (n=72).

### 3.4 Antibiotypes of *S. aureus* from the food samples

A total of 24 antibiotypes were identified in this study (Table 7). The most prevalent antibiotype was P-AML-E-AZM-C that was detected in 9 (12.5%) of the 72 *S. aureus* isolates. More antibiotypes were identified in fruit salad followed by meat-hot-pot.

**Table 7:** Antibiotic-resistant patterns of *S. aureus* isolates from RTE food samples (n=72)

| Number of antibiotics showing resistance | Antibiotype | Number of isolates (%) | Sample origin |
| --- | --- | --- | --- |
| 2 | P-DA | 2 (2.78) | Fruit salad, suya |
|  | E-CN | 1 (1.39) | Meat-hot-pot |
|  | P-AML | 1 (1.39) | Cake |
|  | AML-E | 2 (2.78) | Fruit salad |
| 3 | E-AZM-C | 1 (1.39) | Meat-hot-pot |
|  | P-AML-E | 1 (1.39) | Suya |
|  | P-AML-DA | 3 (4.17) | Rice, Bread |
|  | AML-E-CN | 2 (2.78) | Fruit salad |
|  | P-AML-AZM | 6 (8.33) | Cake, rice, bread |
| 4 | P-E-AZM-C | 1 (1.39) | Meat-hot-pot |
|  | P-AML-E-DA | 6 (8.33) | Meat-hot-pot, rice |
|  | P-E-AZM-CN | 1 (1.39) | Fruit salad |
|  | P-AML-E-CN | 2 (2.78) | Fruit salad |
|  | P-AML-E-AZM | 8 (11.11) | Suya, bread, cake |
|  | P-AML-DA-CN | 1 (1.39) | Meat-hot-pot |
| 5 | P-AML-E-AZM-C | 9 (12.5) | All the RTE foods |
|  | P-AML-E-DA-CN | 5 (6.94) | Fruit salad, rice |
|  | P-AML-CIP-E-CN | 2 (2.78) | Meat-hot-pot |
|  | P-AML-E-AZM-DA | 7 (9.72) | Rice, bread, cake |
|  | P-AML-E-AZM-CN | 1 (1.39) | Fruit salad |
| 6 | P-AML-OX-CIP-E-C | 1 (1.39) | Fruit salad |
|  | P-AML-E-AZM-DA-CN | 6 (8.33) | Rice, suya |
| 7 | P-AML-OX-CIP-E-AZM-C | 2 (2.78) | Meat-hot-pot |
| 9 | P-AML-OX-CIP-E-AZM-DA-CN-C | 1 (1.39) | Bread |
| Total |  | 72 (100) |  |

P, penicillin; AML, amoxicillin, OX, oxacillin; CIP, ciprofloxacin; E, erythromycin; AZM, azithromycin; DA, clindamycin; CN, gentamicin; C, chloramphenicol; RTE, ready-to-eat

## 4. Discussion

The high rate of consumption of RTE foods in Cameroon and especially in the Buea municipality coupled with the scarcity of data on the microbiological quality of these foods is a call for concern. Elsewhere, there are several reports that highlight the role of RTE foods in the emergence or reemergence of microbial threats to health, by providing information on the contamination of RTE foods by bacteria or resistant bacterial species to guide public health and food safety interventions [38]. Staphylococcal food poisoning, caused by the consumption of one or more of the 23 heat-stable enterotoxins and many variants elaborated by coagulase-positive staphylococci and particularly *Staphylococcus aureus*, is a global public health concern [13, 39]. Most food types, including RTE foods, support the growth of staphylococci and are ideal for enterotoxin production [23]. Although staphylococcal food poisoning is usually self-limiting and resolves within 24 to 48 h after onset, occasionally it can be severe and require hospitalization [1]. *Staphylococcus aureus* is ubiquitous in the environment, a commensal in the human body and a notorious zoonotic and major foodborne pathogen that quickly acquires resistance to antibiotics (Kakoullis et al., 2021; Guo et al., 2020). Therefore this study that investigated RTE foods for contamination with coagulase-positive staphylococci with particular focus on *S. aureus* and its antibiotic-resistant profiles, has both public health and food safety relevance.

Of the 420 RTE food samples analyzed in this study, 161 (38.3 %) had detectable levels of CoPS contamination with counts ranging from 2.0-5.81 log_10_CFU/g. Based on the CoPS counts, 37 (8.81%) of the 420 RTE food samples investigated and comprising only of fruit salad and meat-hot-pot, had unsatisfactory or unacceptable quality (Table 4). Our findings corroborate a previous study that analyzed 275 RTE food samples and reported that 25 (9.1%) were inappropriate for consumption according to Turkish microbiological guidelines [40]. Monitoring CoPS count in RTE foods is very helpful for a risk assessment. The number of CoPS per gram or milliliter of RTE food sample is an indication of the microbiological quality of the food sample [20]. In an earlier study that analyzed 60 RTE foods served in long-term care facilities in spain, the highest CoPS count in the samples was 1.9 ± 0.3 log_10_CFU/g, hence all samples had acceptable quality based on United Kingdom guidelines [41]. In a very recent study that investigated 83 RTE food samples, 25% (23/83) of those analyzed were classified as unsatisfactory or potentially hazardous [42].

In this study, fruit salad samples showed high levels (51/70, 72.9 %) of contamination with CoPS (Table 3). High contamination of RTE fruit salads has been reported in previous studies. Brooks [43] reported a higher rate (90 %) of staphylococcal contamination of 20 RTE salad samples analyzed in his study carried out in Calabar, Nigeria. In another recent study that analyzed 36 RTE pineapple slices in Port Harcourt, Nigeria, high levels of bacterial contamination were detected in all the samples and the contamination levels were even higher for samples collected in the evening than for those collected in the morning from the same vendors [44]. The high levels of bacterial contamination of RTE fruits could be multifactorial and include: washing the fruits with poor quality source of water, cross contamination from other fruits washed in the same water many times [45, 46], use of dirty trays or dirty processing utensils like knives and slicing tables, not washing the hands thoroughly and contamination from the air [44].

A total of 12 (17.14%) of the 70 suya samples analyzed in this study had the lowest detectable levels of CoPS and it ranged from 2 - 2.91 log_10_CFU/g (Table 3). While these 12 samples had borderline quality, the remaining 58 (82.86%) samples had satisfactory quality (Table 4). Hence, all the 70 suya samples analyzed in this study were suitable for consumption at the time of sample collection. Falegan et al. [47] in a recent study carried out in Ado-Ekiti Metropolis, Ekiti State, Nigeria, reported comparably higher values of 4.99 - 5.45 log_10_CFU/g for total bacterial counts in the 20 suya samples analyzed and stated that all the samples had unsatisfactory quality. In another earlier study by Egbebi and Seidu [48] carried out in Ado and Akure, South West Nigeria, bacterial counts ranged from 4.48 - 4.60 (in Ado) and 4.48 – 4.93 (in Akure), and these values placed the suya samples in acceptable but not satisfactory range. The low values of CoPS in the suya samples reported in our study could be linked to the fact that these samples were purchased shortly after preparation (i.e freshly prepared) and not much post-processing contamination had occurred.

In this study, majority of the samples of boiled rice (75.71%), bread (61.43%) and cake (70.0%) (Table 4) had satisfactory quality while 24.29% (boiled rice), 38.57% (bread) and 30% (cake) samples had CoPS counts on the high threshold but within microbiologically acceptable limits recommended by regulatory bodies for vended meat and meat products [14, 15]. Our results did not agree with that of Wogu et al. [49] who reported that most of the 48 rice samples from hotels in Benin City, Nigeria examined in their study had unsatisfactory quality. A very recent study that examined 50 rice samples obtained from cafeterias in Akwa Ibom State University in Nigeria reported that most of the rice samples had unsatisfactory quality [50]. The reasons for the differences in the results reported in these studies could be multifactorial ranging from rice species differences, methods of preparation, holding temperature and time and methods of sample analysis.

*Staphylococcus aureus* was identified in 46 (10.95 %) RTE food samples in this study. This overall contamination rate by *S. aureus* reported in our study was higher than the 6.6% (8/120) reported for RTE foods in Egypt [51], 4.31% (6/139) in Libya [52] and 6.4% (48/750) in Istanbul in Turkey [53] but lower than the 29.1% (155/532) in Cambodia [12], 34% (68/200) in Bolgatanga municipality of Ghana [54], 52.8% (56/106) in Putrajaya, Malaysia and 12.55% (69/550) in China [25]. The difference in the prevalence rates reported in these studies could be due to diagnostic method used; while some studies identified *S. aureus* using phenotypic methods, others relied on the more sensitive molecular method.

A total of 72 *S. aureus* isolates comprising 52.78% from fruit salad, 16.67% from meat-hot-pot, 12.5% from boiled rice, 9.72% from suya, 5.56% from bread and 4.17% from cake (Table 5) were recovered from the 46 S. aureus-positive samples in this study. Since *S. aureus* is heat-labile and also readily destroyed by most sanitizing agents, its presence in RTE foods may be an indication of poor handling and sanitation [38]. Elsewhere, contamination of RTE foods has been linked to the preparation by small-scale local producers without quality control, re-use of improperly washed dishes, open-air distribution environment and holding temperatures during distribution [55]. We investigated the *S. aureus* isolates for the presence of the classical enterotoxin genes and none was found to possess any of the genes. In a similar study that investigated 84 *S. aureus* isolates for the presence of the classical enterotoxin genes, only 2 (2.38%) of the isolates possessed the enterotoxin B gene while the rest of 82 isolates were negative for other enterotoxins genes [23]. In another study, a high prevalence (58.1%, 18/31) of classical staphylococcal enterotoxin genes was identified in meat samples in Zanjan, Iran [56]. Although a positive PCR confirms the presence of enterotoxins, the negative PCR in our study does not point to the absence of the corresponding operon because there is always a possibility of mutation at the level of the corresponding gene. Furthermore, the differences between our results and those reported elsewhere could also be due to other factors like sample source and geographical origin [23].

The increasing reports that resistant strains of *S. aureus* are present in various RTE foods in different countries [55] and the notorious resistance of this bacterium to several commonly used antibiotics [57] warranted antibiotic susceptibility testing for all 72 *S. aureus* isolates identified in this study. All the *S. aureus* isolates showed 100% sensitivity to vancomycin and ofloxacin followed by ciprofloxacin (91.67%) (Table 6). These results agree with that of Sina et al. [58] in a study carried out in Cotonou Benin and that of Wang et al. [59], who reported that all *S. aureus* isolates from food in Shaanxi Province, China were sensitive to vancomycin. This high sensitivity of the *S. aureus* isolates has great therapeutic relevance because these antibiotics can be used to treat infections caused by this bacterium in our resource poor community. High *S. aureus* antibiotic resistance was observed for penicillin (93.06%) followed by amoxicillin and erythromycin and our results are comparable to that reported by Tsehayneh et al. [60]. Forty-five (62.5%) *S. aureus* isolates in this study showed multidrug resistance and this was lower than 98% (54 out of 55 *S. aureus* isolates) recently reported by Tsehayneh et al. [60]. This difference in resistance reported may be due to the extent of usage of antibiotics in a locality and possibly the type of samples analyzed [55].

In this study, resistance to oxacillin was used to identify methicillin-resistant *S. aureus* (MRSA) strains. A total of four (5.56%) of the 72 *S. aureus* investigated were MRSA and all possessed the *mec*A gene. Saad et al. [51] reported that two (25%) of the eight *S. aureus* isolates identified in their study were MRSA, Yang et al. [25] reported 10.14% (7/69) prevalence of MRSA out of which six were *mec*A-positive, Naas et al. [52] reported only one (16.67%) of six *S. aureus* isolates as MRSA but did not screen for the presence of the *mec*A gene. All MRSA strains in this study were sensitive to vancomycin and as expected, but resistant to penicillin and other antibiotics. MRSA is a pathogen of global concern and is placed in the WHO priority pathogen list [61] because infections caused by it are extremely difficult to treat. Thus, understanding the epidemiology of this pathogen and its sources in the environment are relevant towards effective prevention and control, especially at this time when the antibiotic pipeline is nearly empty [61]. A total of 24 antibiotypes were identified among the 72 *S. aureus* isolates in this study and they did not show any clustering with the origin of the *S. aureus* isolates.

## 5. Conclusion

This study presents very important findings in terms of public health and food safety because these RTE foods are consumed without further heat treatment. The potential role of these RTE foods in the spread of CoNS and *S. aureus* cannot be ignored. The contamination of these foods with CoNS, S. *aureus* and even MRSA poses a potential risk to consumers. To combat the spread of antibiotic-resistant *S. aureus* and particularly MRSA, it is necessary and urgent to identify all possible sources and potential pathways by which these pathogens spread. Moreover, our study also provides insight into the antibiotypes of S. aureus circulating in the study area.

## Acknowledgements

The authors are grateful to the Laboratory for Emerging Infectious Diseases, University of Buea, for providing equipment to accomplish this work.

## Author contributions

Conceptualization: Seraphine Nkie Esemu, Lucy M. Ndip

Data curation: Seraphine Nkie Esemu, Nene Kaah Keneh, Jerome Achah Kfusi

Formal analysis: Seraphine Nkie Esemu, Nene Kaah Keneh, Sally Tabe Njoh, Jerome Achah Kfusi

Investigation: Seraphine Nkie Esemu, Nene Kaah Keneh, Sally Tabe Njoh, Jerome Achah Kfusi

Methodology: Seraphine Nkie Esemu, Lucy M. Ndip, Achiangia Patrick Njukeng

Resources: Seraphine Nkie Esemu, Sally Tabe Njoh, Lucy M. Ndip

Supervision: Seraphine Nkie Esemu, Achiangia Patrick Njukeng

Validation: Lucy M. Ndip Achiangia Patrick Njukeng

Writing – original draft: Seraphine Nkie Esemu

Writing – reviewing and editing: Lucy M. Ndip Achiangia Patrick Njukeng

